# Evolutionary forces on different flavors of intrinsic disorder in the human proteome

**DOI:** 10.1101/653063

**Authors:** Sergio Forcelloni, Andrea Giansanti

**Author notes:** Corresponding Author: Sergio Forcelloni, Sapienza University of Rome, Department of Physics, P.le A. Moro 5, 00185 Roma, Italy.

## Abstract

In this study, we perform a systematic analysis of evolutionary forces (i.e., mutational bias and natural selection) that shape the codon usage bias of human genes encoding for different structural and functional variants of proteins. Well-structured proteins are expected to be more under control by natural selection than intrinsically disordered proteins because one or few mutations (even synonymous) in the genes can result in a protein that no longer folds correctly. On the contrary, intrinsically disordered proteins are generally thought to evolve more rapidly than well-folded proteins, primarily attributed to relaxed purifying natural selection due to the lack of structural constraints. Using different genetic tools, we find compelling evidence that intrinsically disordered proteins are the variant of human proteins on which both mutational bias and natural selection act more effectively, corroborating their essential role for evolutionary adaptability and protein evolvability. We speculate that intrinsically disordered proteins have a high tolerance to mutations (both neutral and adaptive) but also a selective propensity to preserve their structural disorder, i.e., flexibility and conformational dynamics under physiological conditions. Additionally, we confirm not only that intrinsically disordered proteins are preferentially encoded by GC-rich genes, but also that they are characterized by the highest fraction of CpG-sites in the sequences, implying a higher susceptibility to methylation resulting in C-T transition mutations. Our results provide new insight about protein evolution and human genetic diseases identifying intrinsically disordered proteins as reservoirs for evolutionary innovations.

## 1. Introduction

The genetic code is redundant, with more than one codon encoding for the same amino acid. Increasing evidence has shown that synonymous codons are not used with equal frequencies, a phenomenon called *codon usage bias (CUB)* (Clarke, 1970; Ikemura, 1985; Plotkin and Kudla, 2011; Shabalina et al., 2013; Hanson and Coller, 2018).

The most accepted theory to explain the origin of CUB is the *selection-mutation-drift theory*, according to which *natural selection* and *mutational bias* are the two leading evolutionary forces shaping CUB (Bulmer, 1991).

Natural selection can influence codon choice of specific genes or even specific codon positions that require a fine control (e.g., at the level of translation and co-translational protein folding) (Roth et al., 2002; Hershberg and Petrov, 2008). That is exemplified in highly expressed genes, where synonymous codon usage patterns are shaped by selection for specific codons that are more efficiently and/or accurately translated by the most abundant tRNAs (dos Reis et al., 2004). On the contrary, mutational bias tends to accumulate asymmetrically certain types of mutations on the whole genome of an organism. This is caused by underlying molecular mechanisms, including but not limited to: methylation of CpG dinucleotide to form 5-methylcytosine and subsequent deamination that results in C-T substitution (Kaufmann and Paules, 1996), chemical decay of nucleotide bases (Kaufmann and Paules, 1996), transcription-associated mutational biases (Green et al., 2003; Comeron, 2004; Cui et al., 2012), non-random error during DNA replication (Lobry, 1996, Cui et al., 2012), biased gene conversion (Eyre-Walker, 1993; Galtier et al. 2001), and non-uniform DNA repair (Roth et al., 2012).

Depending on the effect of the mutation on the fitness of the organism, we can distinguish three broad categories of mutations (Fordsyke, 2016). I) If the fitness is decreased by a mutation (*deleterious mutation*), then the selection of the organism with such mutation is negative (*negative selection*). II) If the fitness is increased by a mutation (*advantageous mutation*), then the selection of the organism with such mutation is positive (*positive selection*). III) If the effect of the mutation on the fitness is weak (*neutral mutation* or *nearly neutral mutation*), such mutation is fixed without being affected by natural selection (Kimura, 1968, Kimura, 1991, Ohta, 1992). Because many sites within genes can undergo synonymous substitutions which do not change the protein sequence encoded, the nucleotide composition of these silent sites is typically considered as a reflection of the mutation patterns (Sharp et al., 1995).

It has been debated for quite a long time about whether selection does not exist or it is too weak to be detectable in human genomes (Ma et al., 2014), where sequence nucleotide patterns are embedded into compartmentalized large regions (>>200 kilobases) of low or high GC-content (the so-called *isochores*) (Bernardi et al., 1985). Although many aspects of the evolution of isochores have long been investigated (Bernardi et al., 1985; Galtier and Mouchiroud, 1998; Sueoka and Kawanishi, 2000; Eyre-Walker and Hurst, 2001; Lercher et al., 2003), it is not entirely clear whether the isochore maintenance is determined by mutational bias or selection or both (Eyre-Walker and Hurst, 2001; Li, 2013). The most likely scenario is that GC-content is mainly determined by genome-wide mutational processes rather than by selective forces acting specifically on coding regions (Sueoka, 1962; Wolfe et al., 1989; Chen et al., 2004; Hershberg and Petrov, 2008; Plotkin and Kudla, 2011; Romiguier and Roux, 2017).

Some evidence has shown the action of selection in the human genome (Lavner and Kotlar, 2005; Comeron, 2004) and, particularly, on housekeeping (Ma et al., 2014) and highly expressed genes (Urrutia and Hurst, 2003). However, it remains not well explored whether evolutionary pressures act differently on human genes depending on the structural properties of the encoded proteins (the aim of this study). Intrinsically disordered proteins are generally thought to evolve more rapidly than globular proteins, due to the lack of structural constraints (Brown et al., 2002; Brown et al., 2010; Brown et al., 2011; Schlessinger et al., 2011; Xue et al., 2013). Accumulating evidence suggests that relaxed structural constraints provide an advantage when accommodating multiple overlapping functions in coding regions (collectively referred to as synonymous constraint elements) (Castillo et al., 2014; Pancsa and Tompa, 2016). Additionally, recent studies have found a higher degree of positive selection on proteins with intrinsically disordered regions (Nillson et al., 2011; Afanasyeva et al., 2018).

Herein, using different genetic tools (ENC-plot (Wright, 1990), Neutrality plot (Sueoka, 1988), PR2-plot (Sueoka, 1995)), we performed a systematic analysis of the evolutionary pressures (i.e., natural selection and mutational bias) that shape CUB of human genes encoding for different structural variants of proteins. We used the operational classification of the human proteome by Deiana et al. (Deiana et al., 2019) to distinguish three variants of human proteins characterized by different structural properties and different functional spectra. We thus considered: i) ordered proteins (ORDPs), ii) structured proteins with intrinsically disordered protein regions (IDPRs), and iii) intrinsically disordered proteins (IDPs). Our results strongly suggest that intrinsically disordered proteins are not only affected by a basic mutational bias, but they also show a peculiar influence of natural selection as compared to the rest of human proteome. In line with that, we find evidence about the important role of intrinsically disordered proteins for evolvability and adaptability of organisms or cells in which they occur.

## 2. Materials and Methods

### 2.1 Data sources

Human proteome was downloaded from the UniProtKB/SwissProt database (manually annotated and reviewed section - https://www.uniprot.org/uniprot/?query=reviewed:yes) (UniProt Consortium, 2017). Human Coding DNA Sequences (CDSs) were retrieved by Ensembl Genome Browser 94 (https://www.ensembl.org/index.html) (Zerbino et al., 2018). Only genes with UniProtKB/SwissProt ID have been included to make sure we only consider coding sequences for proteins. We consider only CDSs that start with the start codon (AUG), end with a stop codon (UAG, UAA, or UGA), and have a multiple length of three. Each CDS was translated in the corresponding amino acid sequence and then we filter all sequences that do not have a complete correspondence with a protein sequence in UniProtKB/SwissProt. Incomplete and duplicated gene, sequences with internal gaps, unidentified nucleotides were removed from the analysis. A list of 18214 human CDSs was generated; the coverage of the human proteome in SwissProt (reviewed) is 90%.

### 2.2 Disorder prediction

We identified disordered residues in the protein sequences using MobiDB3.0 (http://mobidb.bio.unipd.it) (Piovesan et al., 2018), a consensus database that combines experimental data (especially from X-ray crystallography, NMR, and cryo-EM), manually curated data, and disorder predictions based on various methods.

### 2.3 Classification of disordered proteins in the human proteome

We partitioned the human proteome into variants of disorder following the operational classification by Deiana et al. (Deiana et al., 2019). In line with that, we distinguish three variants of human proteins having different structural and functional properties: i) ordered proteins (ORDPs), ii) structured proteins with intrinsically disordered protein regions (IDPRs), and iii) intrinsically disordered proteins (IDPs). Generally speaking, ORDPs are proteins with a limited number of disordered residues and the absence of disordered domains. IDPRs, unlike ORDPs, are proteins with at least one long disordered domain accommodated in globally folded structures. IDPs are proteins with a relevant fraction of disordered residues in the sequence (more than 30%).

### 2.4 GC-content

The GC-content of a gene is the percentage of guanine and cytosine bases with respect to the total length of that gene. Similarly, it is possible to define the GC-content in the first (GC1), second (GC2), and third (GC3) codon positions, as follows:

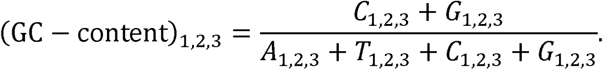

### 2.5 The Effective Number of Codons

We calculated the effective number of codon (ENC) to estimate the extent of the codon usage bias of human genes encoding for ORDPs, IDPRs, and IDPs. The values of ENC range from 20 (when just one codon is used for each amino acid) to 61 (when all synonymous codons are equally used for each amino acid) (Wright, 1990). Thus, the smaller is the ENC value the larger is the extent of codon preference in a gene. The ENC values of all individual human genes (>600 bp) were calculated by using the improved implementation by Sun et al. (Sun et al., 2012) as follows. The codon table was separated in synonymous codon families. The 6-fold codon families (Leu, Ser, and Arg) were divided into 2-fold and 4-fold codon families. For each coding sequence, we quantify Fα, defined for each synonymous codon family α as:

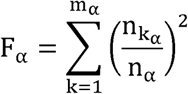

where m_α_ is the number of codons in the codon family α, n_iα_ with *i* = 1,2,…,*m*_*α*_ is the number of occurrences of codon *i* of the codon family α in the coding sequence, and 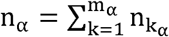. Finally, the gene-specific ENC is defined as:

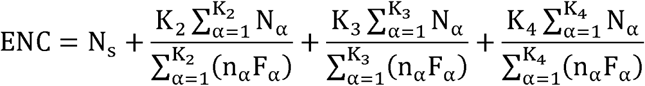

where N_s_ is the number of codon families with a single codon (i.e., Met and Trp codon families) and K_m_ is the number of families with degeneracy m.

### 2.6 ENC-plot

An ENC-plot analysis (Wright, 1990) was performed to estimate the relative contributions of mutational bias and natural selection in shaping CUB of human genes encoding for ORDPs, IDPRs, and IDPs. The ENC-plot is a plot in which the ENC is the ordinate and the GC3 is the abscissa. Depending on the action of mutational bias and natural selection, different cases are discernable. If a gene is not subject to selection, a clear relationship is expected between ENC and GC3:

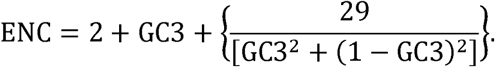

Genes for which the codon choice is only constrained by mutational bias are expected to lie on or just below the Wright’s theoretical curve. Alternatively, if a particular gene is subject to selection, it will fall below the Wright’s theoretical curve. In this case, the vertical distance between the point and the theoretical curve provides an estimation of the relative extent to which natural selection and mutational bias affect CUB.

### 2.7 Neutrality plot

We used a neutrality plot analysis (Sueoka, 1988; Sueoka, 1999) to compare the relative neutrality of the first and second codon positions compared to that of third codon position (generally a silent site) of genes encoding for ORDPs, IDPRs, and IDPs. In this analysis, the GC1 or GC2 values (ordinate) are plotted against the GC3 values (abscissa), and each gene is represented as a single point on this plane. For each variant of disorder (ORDPs, IDPRs, and IDPs), we separately performed a Spearman correlation analysis between GC1 and GC2 with the GC3. If the correlation between GC1/2 and GC3 is statistically significant, the slope of the regression line provides a measure of the relative neutrality of the first/second codon position with respect to the third one (Sueoka, 1999). In particular, if the first or second codon position is as neutral as the third codon position, then the corresponding data points should be distributed along the bisector (slope of unity). On the other hand, if the first or second codon position is completely non-neutral, then the corresponding data points should be on the abscissa (slope of zero). Thus, a slope smaller than unity indicates that the extent of neutrality in first or second codon positions is less than that of third codon position.

### 2.8 Parity Rule 2 (PR2) plot

A Parity Rule 2 (PR2) plot analysis was performed to assess the relative contributions of mutational bias and natural selection on the CUB of genes encoding for ORDPs, IDPRs, and IDPs. In this plot, the GC-bias [G3/(C3+G3)|_4_)] and the AT-bias [A3/(A3+T3)|_4_] at the third codon position of the four-codon amino acids of individual genes are plotted as the abscissa and the ordinate respectively (Sueoka, 1995; Sueoka, 1999). When there is no bias between the effects of mutational bias and natural selection, data points should distribute around the center where A3=T3 and G3=C3 (PR2). Alternatively, selective forces rather than mutational biases during evolution are expected to lead to deviations from the PR2 (Sueoka, 1995). In this case, data points should move away from the center, and the distance from it provides a measure of the relative effects of natural selection and mutational bias on the base composition and codon usage of genes.

### 2.9 Quantification of CUB not due to the background compositional bias

The analyses based on ENC-, Neutrality-, and PR2-plot only allow estimating the *relative (but not the net)* contributions of natural selection and mutational bias in shaping CUB of genes. The isochore structure of the human genome underlines the need to correct for the effect of background (di)nucleotides composition when attempting to detect the net contribution of natural selection on CUB (Chamary et al., 2006). A statistical correction (ENC’) was developed to take into account the wide variability of the GC-content (Novembre, 2002). Although this is an improvement of Wright’s original method, ENC’ has different limitations (Fuglsang 2006; Belalov and Lukashev, 2013): i) it is dependent from the length of the coding sequences; ii) it provides inaccurate estimates for short sequences (<600 bp); iii) it does not explicitly account for nucleotide and dinucleotide compositional bias that may be high in the human genomes. Moreover, it is possible that by using this method we eliminate some of the signals we expect to find (Hershberg and Petrov, 2009). For these reasons, we followed the procedure by Belalov and Lukashev (Belalov and Lukashev, 2013), who developed different shuffling techniques of the coding sequences to filter all effects resulting from background (di)nucleotides compositions. Briefly, each technique takes into account a positional (di)nucleotide contents on specific codon positions (e.g., the nucleotide content at third codon position) and shuffles each coding sequence with respect to such codon positions with the constraint to preserve the amino acid sequence. Because the positional (di)nucleotide composition is maintained by definition, this procedure enables to evaluate the extent of CUB that is not due to the effect of background (di)nucleotides bias, which itself is likely due to the mutational bias. Detailed descriptions of the shuffling techniques are reported in the article by Belalov and Lukashev (Belalov and Lukashev, 2013) and summarized below. *I) N*_*3*_ *correction* shuffled third codon positions while preserving the base composition at the third codon position and the amino acid sequence. *II) dN23* (*dN31*) *correction* shuffled dinucleotides between codon positions 2-3 (3-1) while preserving the dinucleotide composition at the codon position 2-3 (3-1) and the amino acid sequence.

### 2.10 CpG dinucleotide relative abundances

The CpG dinucleotide relative abundance is defined as:

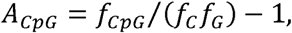

where *f*_*CpG*_ is the frequency of CpG dinucleotide in the sequence and *fC*(*f*_*G*_) is the frequency of cytosine (guanine) in the variant of disorder. Generally speaking, *A*_*CpG*_ quantifies the abundance of CpG sites relative to what would be expected from the base composition of the variant. By calculating *A*_*CpG*_ for each variant of proteins (ORDPs, IDPRs, and IDPs), we can have three possible cases: i) if *A*_*CpG*_ = 0, then the frequency of CpG sites is coherent with the base composition of the variant; ii) if *A*_*CpG*_ < 0, then the CpG sites are underrepresented in the variant; iii) if *A*_*CpG*_ > 0, then the CpG sites are overrepresented in the variant.

## 3. Results

### 3.1 ENC-plot of the three variants of disorder

In Fig. 1, we show separately the ENC-plots obtained for the three variants of proteins (ORDPs, IDPRs, and IDPs), together with best-fit curves obtained through a nonlinear fit of Wright’s shape, with parameters a, b, c, and d:

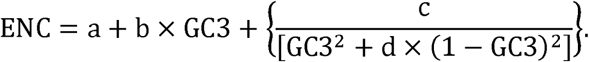

**Fig. 1:**
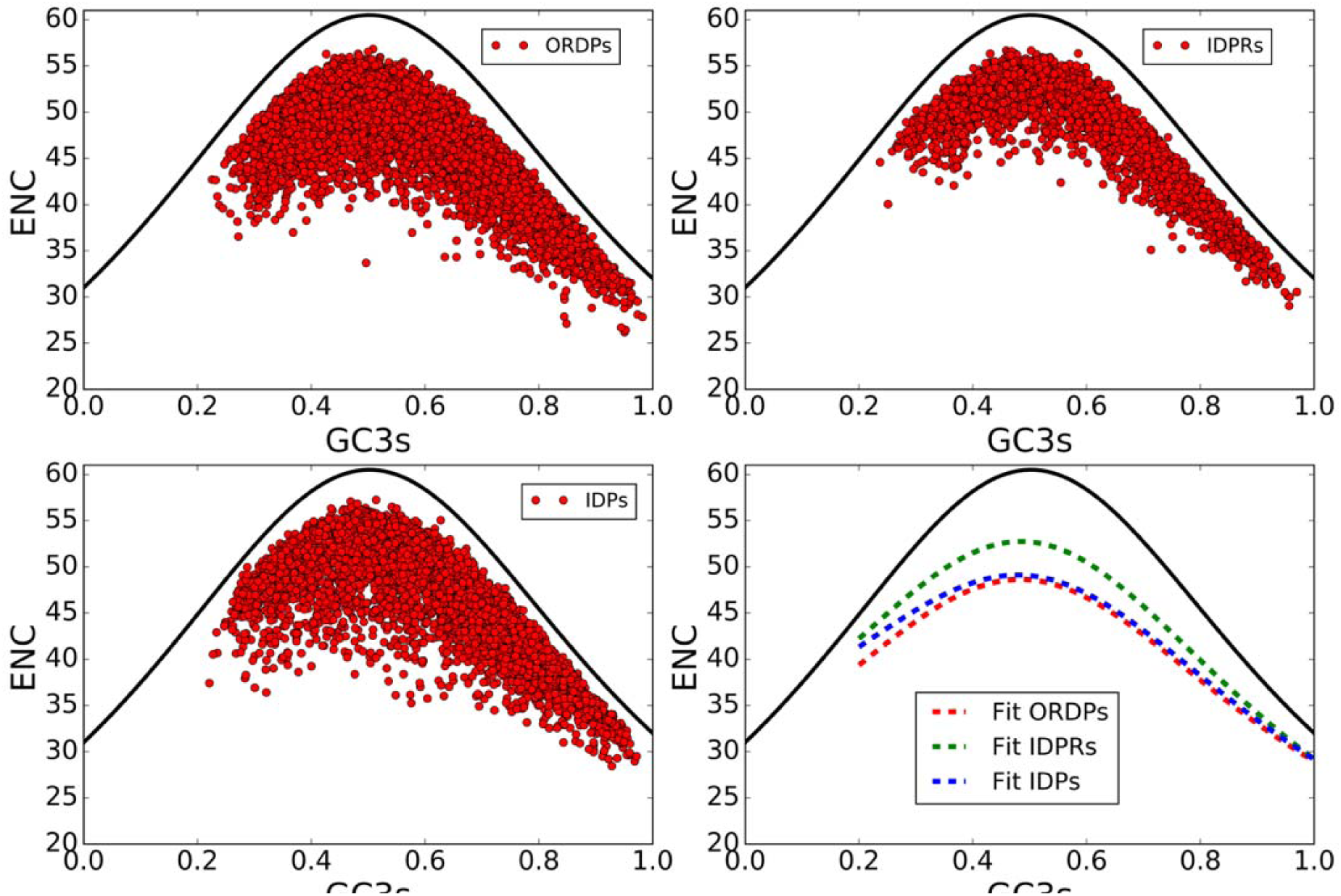
ENC-plots of ORDPs, IDPRs, and IDPs. The solid black lines in all panels are plots of Wright’s theoretical curve (see Materials and Methods) corresponding to the case of a CUB merely due to mutational bias (no selective pressure). In the bottom-right panel dashed lines are non-linear fits of Wright’s theoretical shapes to the experimental data. The best-fit values of the parameters are: ORDPs: a=10.58651558, b=-0.03584713, c=18.41801672, d= 0.93760164; IDPRs: a=9.7618343, b=-2.19167042, c=21.62987183, d=0.96492965; IDPs: a=16.1986867, b=-4.34617166, c=17.31627503, d=0.97792585.

In all the three cases the shape of the experimental distributions follows the general shape of Wright’s theoretical curve, indicating that CUB in all variants of human proteins reflects the GC3 variation, which itself is mainly determined by mutational bias. Nevertheless, it is worth noting that all distributions lie below the theoretical curve, an indication that natural selection also plays a non-negligible role in the codon choice in all the variants. ORDPs and IDPs data points are more scattered below the theoretical curve than those of IDPRs, implying that in the latter the codon usage is under more strict control by GC3. Looking at the best-fit curves in the bottom-right panel, ORDPs and IDPs appear to be subjected on average to a similar balance between mutational bias and natural selection. Conversely, the best-fit curve of IDPRs is closer to Wright’s theoretical curve, meaning that mutational bias affects more markedly the CUB of this variant. All variants of proteins show a displacement towards higher values of GC3-content, which is probably the result of mosaic structure of the human genome (Bernardi et al., 1985).

### 3.2 Parity Rule 2 (PR2) plot of the three variants of disorder

Human genes encoding for ORDPs, IDPRs, and IDPs were plotted on the PR2-plot (Fig. 2). All distributions are located around the center, confirming that mutational bias is the predominant factor shaping CUB of human genes. However, by looking more closely the positions of centroids (represented by stars in the PR2-plot), slightly different PR2 violation pattern can be observed, indicating a differential balance between the influences of mutational bias and natural selection on CUB of genes encoding for the variants of disorder. In particular, the pyrimidines (C and T) seem to be used on average more frequently than the purines (A and G) at the synonymous sites in all variants, especially in ORDPs.

**Fig. 2:**
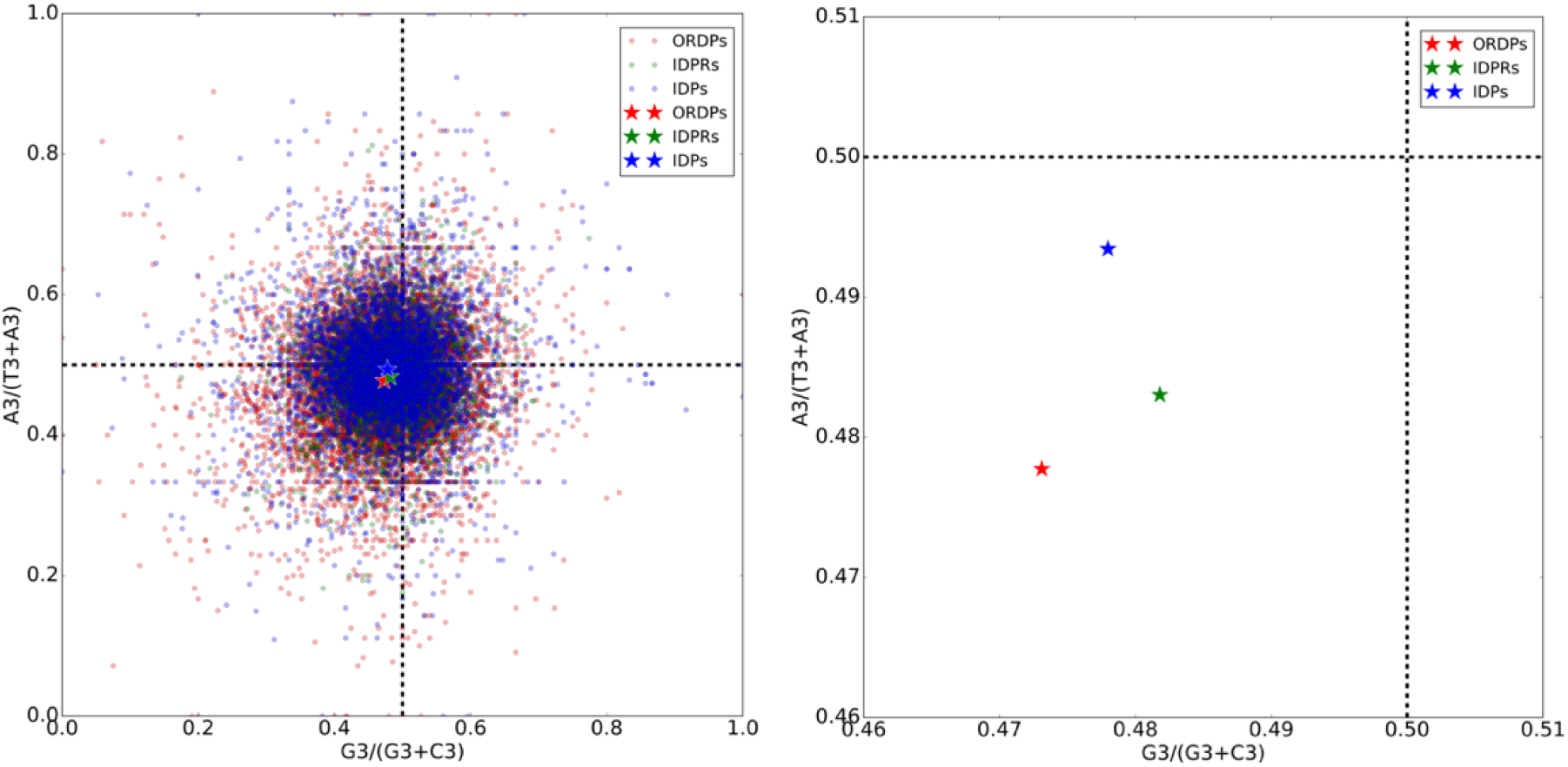
PR2-plot of ORDPs (red), IDPRs (green), and IDPs (blue). Deviations from PR2 in A-T bases of the third codon position are shown as A3/(A3+T3)_|4-fold_ (ordinate) and those in C-G bases are shown as G3/(C3+G3)_|4-fold_ (abscissa) for the eight four-codon family amino acids. The centroids of distributions (stars) are plotted on the magnified PR2-plot on the right, error bars are standard deviations in the mean. From the positions of the centroids, it appears that C and T are more favored than G and A in all variants of human proteins.

In this framework, however, the position of the centroids is not useful to quantify the relative extent of natural selection and mutational bias, because opposite deviations from PR2 for proteins in the same variant are mostly canceled out when they are combined. Thus, for each protein, we calculated the relative distance (*r*) of its point in the PR2-plot from the center:

corresponds to the case of perfect balance between mutational bias and natural selection. On the contrary, if mutational bias and natural selection give different contributions in shaping CUB of a gene then its point on the PR2-plot moves away from the center (). In Fig. 2, we show the distributions of these distances of individual genes in each variant of disorder (ORDPs, IDPRs, and IDPs). The difference between the three *r*-distributions were identified as highly significant (Kruskal-Wallis H-test and Mann-Whitney U-test with Bonferroni correction,). The PR2 violation appears in this order: IDPRs, IDPs, ORDPs, which is consistent with the distances between the curve-fit of the variants and Wright’s theoretical curve in the ENC-plot (Fig. 1). Interestingly, the extent of biases from PR2 is more pronounced in ORDPs and IDPs than in IDPRs, suggesting the presence of a selective pressure slightly higher for ORDPs and IDPs than for IDPRs (Fig. 3).

**Fig. 3:**
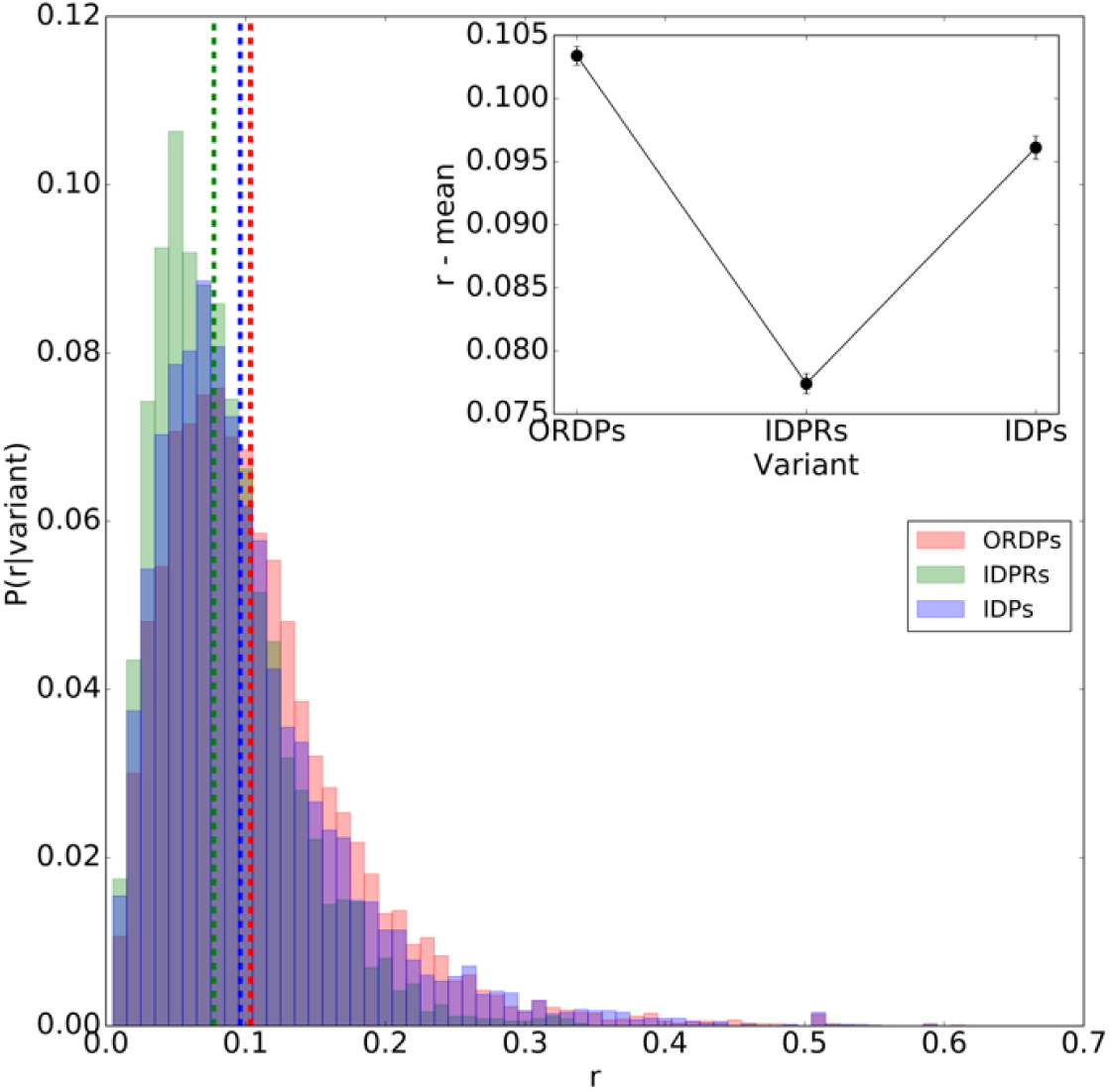
Distributions of the distances of individual genes in ORDPs (red), IDPRs (green), and IDPs (blue) from the center of the PR2-plot. The vertical colored lines represent the average values of distributions. Starting from the origin (), the variants appear in this order: IDPRs, IDPs, ORDPs, meaning that mutational bias and selective pressure are differently balanced in IDPRs, than are in ORDPs and IDPs. In the inset, we show in detail the mean values of distributions for each variant of disorder. The difference between the *r*-distributions were identified as highly significant (Kruskal-Wallis H-test and Mann-Whitney U-test with Bonferroni correction for multiple comparisons, p-value<0.00001).

### 3.3 Neutrality plot of the three variants of disorder

A neutrality plot analysis was performed to estimate the relative neutrality of the first and second codon positions (in general no silent sites), assuming the third codon position (in general silent site) as neutral. In Fig. 4, we report the neutrality plots obtained for ORDPs, IDPRs, and IDPs, together with the best-fit lines and the slopes associated with them. The rationale to understand the results below is that the smaller is the slope of the regression line, the less neutral is the first or the second codon positions with respect to the third one. Looking at Fig. 4, different considerations can be made. Firstly, all correlation are highly significant (Spearman correlation analysis,). Secondly, the wide dispersion in GC3 (from 20% to 100%) reflects the base composition of the local DNA region, which itself is probably the result of variation in mutational bias among chromosomal regions (isochores) (Bernardi et al., 1985). Thirdly, the low values of slopes of the regression lines reveal the strong action of selective pressure on the first two codon positions in all variants of disorder. Fourthly, both first and second codon positions appear to be more neutral in genes encoding for IDP than in genes encoding for ORDPs and IDPRs. Finally, it is worth noting that the slopes in GC1 vs. GC3 plot are systematically steeper than those obtained for GC2 vs. GC3 plot, reflecting the crucial role of the second codon position in determining the chemical-physical properties of the encoded amino acid.

**Fig. 4:**
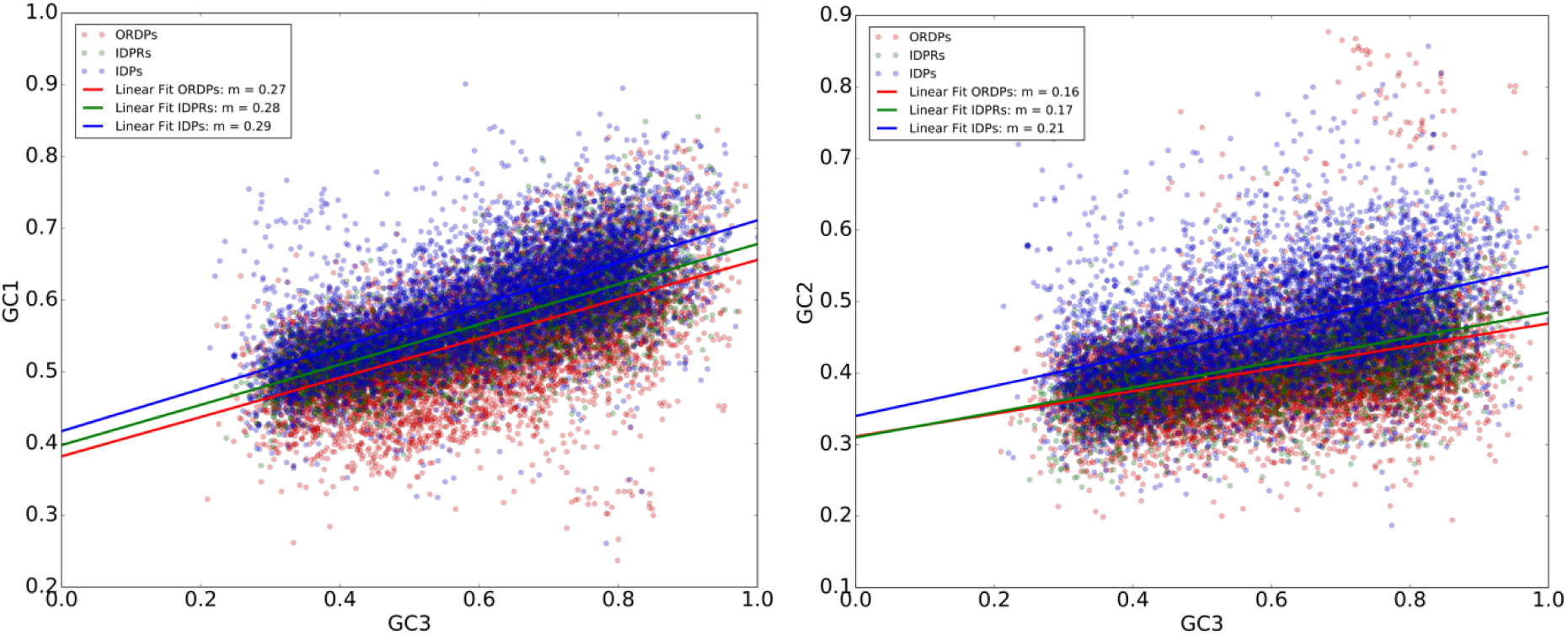
Neutrality plot of ORDPs (red), IDPRs (green), and IDPs (blue). On the left, GC1 vs GC3; on the right, GC2 vs GC3. All linear regressions are characterized by a strong, significant correlation (Spearman correlation analysis,). For each data points, we report the best-fit line and the associated slope of the linear regression in the legend. The slopes (m) of the linear regressions are: GC1 Vs. GC3: ORDPs: m = 0.27, IDPRs: m = 0.28, IDPs: m = 0.29; GC2 Vs. GC3: ORDPs: m = 0.16, IDPRs: m = 0.17, IDPs: m = 0.21.

### 3.4 Variability of the GC-content among variants of proteins

In human as well as in other mammals, CUB mainly depends on the GC-content of the genomic regions (isochores) where the genes reside (Chamary et al., 2006). Although the causes of isochoric structure remain unclear, many evidence indicates that the isochoric structure is related to several genomic features that are functionally relevant, for example, gene density, methylation rate, recombination rate, and expression levels (Eyre-Walker and Hurst, 2001; Kudla et al., 2006, Li 2013; Romiguier and Roux, 2017). For these reasons, we examined separately GC-content of genes encoding for ORDPs, IDPRs, and IDPs. The form of distributions differentiates ORDPs, IDPRs, and IDPs (Fig. 5): i) genes encoding for ORDPs show a unimodal peak for GC-content < 0.5; ii) genes encoding for IDPRs show a bimodal peak; iii) genes encoding for IDPs show a unimodal peak GC-content > 0.5. Looking at the average values of distributions (vertical red lines), we observed a shift of them towards higher values of the GC-content passing from ORDPs, IDPRs, up to IDPs (that is, increasing the percentage of disordered residues) (Fig. 5). This observation suggests that the high GC-content is maintained preferentially in the disordered protein regions.

**Fig. 5:**
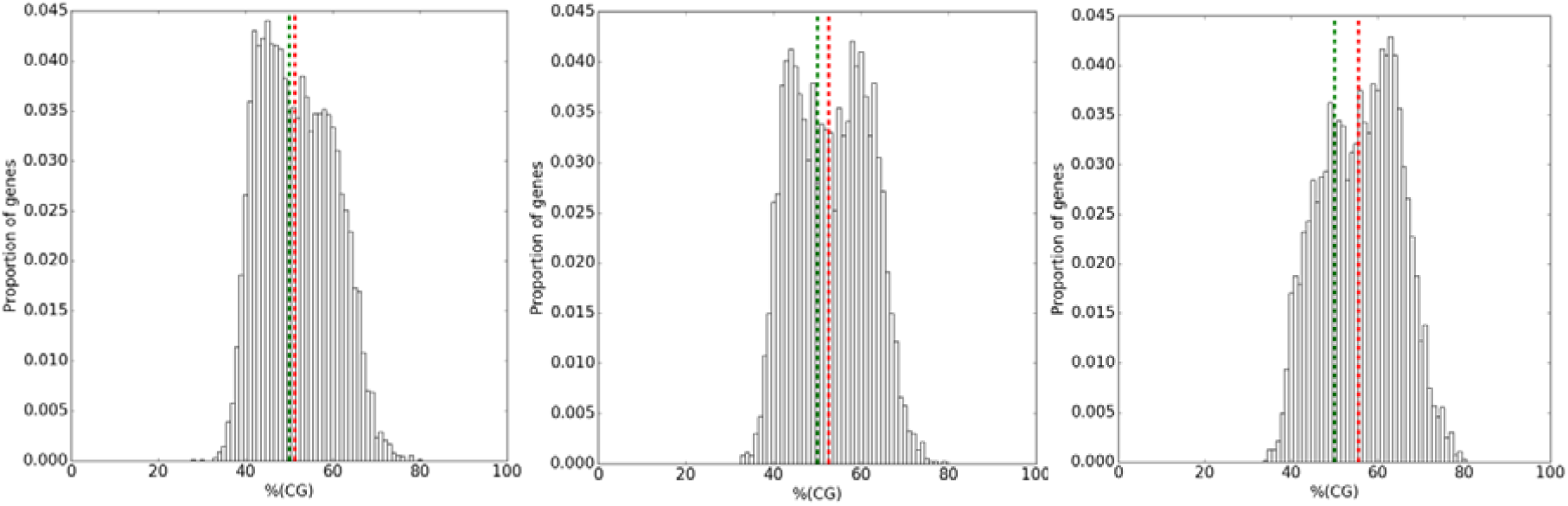
Histograms of GC-content of genes encoding for ORDPs (left), IDPRs (center), and IDPs (right). The vertical red lines represent the average values of distributions; the vertical green lines indicate where.

In support of that, we analyzed the average GC-content for each variant of disorder and for different percentiles of disordered residues in IDPs (Fig. S1). In line with Peng et al. (Peng et al., 2016), an increasing trend is observed confirming the positive correlation between the GC-content of the genes and the percentage of disordered residues in the corresponding protein sequences.

### 3.5 Estimation of the effect of background compositional bias on CUB of the variants

The mutational explanation posits that CUB mainly arises from biases in (di)nucleotides composition along the human chromosomes (e.g., in GC-content) that are caused by molecular mechanisms that favor unidirectionally specific types of mutations (Roth et al., 2012; Li et al., 2015). For this reason, quantifying the net contribution of natural selection on CUB can be difficult and requires a correction for the effect of background (di)nucleotide composition (Chamary et al., 2006).

The standard procedure to estimate the impact of (di)nucleotide compositional bias on CUB is shuffling codons in the real coding sequence while preserving the factor under study (i.e., a specific positional nucleotide or dinucleotide content) and the corresponding amino acid sequence (Belalov and Lukashev, 2013). Thus, for each real codon sequence, we can have the corresponding shuffled version that satisfies the constraints on which it is possible to calculate the corresponding ENC. Depending on the values of ENC associated with the real and shuffled codon sequence, two cases are possible. The first one, if the ENC of the shuffled sequence is equal to the ENC of the real coding sequence, then the constraint fully explains the CUB (i.e., with that (di)nucleotide composition, it is not possible to have ENC higher than the original one). The second one, if the ENC of the shuffled sequence is higher than ENC of the real coding sequence, then the (di)nucleotide compositional bias used as a constraint for the shuffling algorithm does not fully explain the extent of CUB.

Firstly, we analyzed the impact of nucleotide composition in the third codon position (N_3_) on the CUB of the variants. In this case, the ENC values obtained for the shuffled sequences were on average higher than the ENC values of the real coding sequences by more than 3 in all variants of disorder, indicating the presence of an unexplained residual bias, but at the same time were below 61, showing the non-negligible effect of N_3_ on CUB (Fig. 7).

**Fig. 7:**
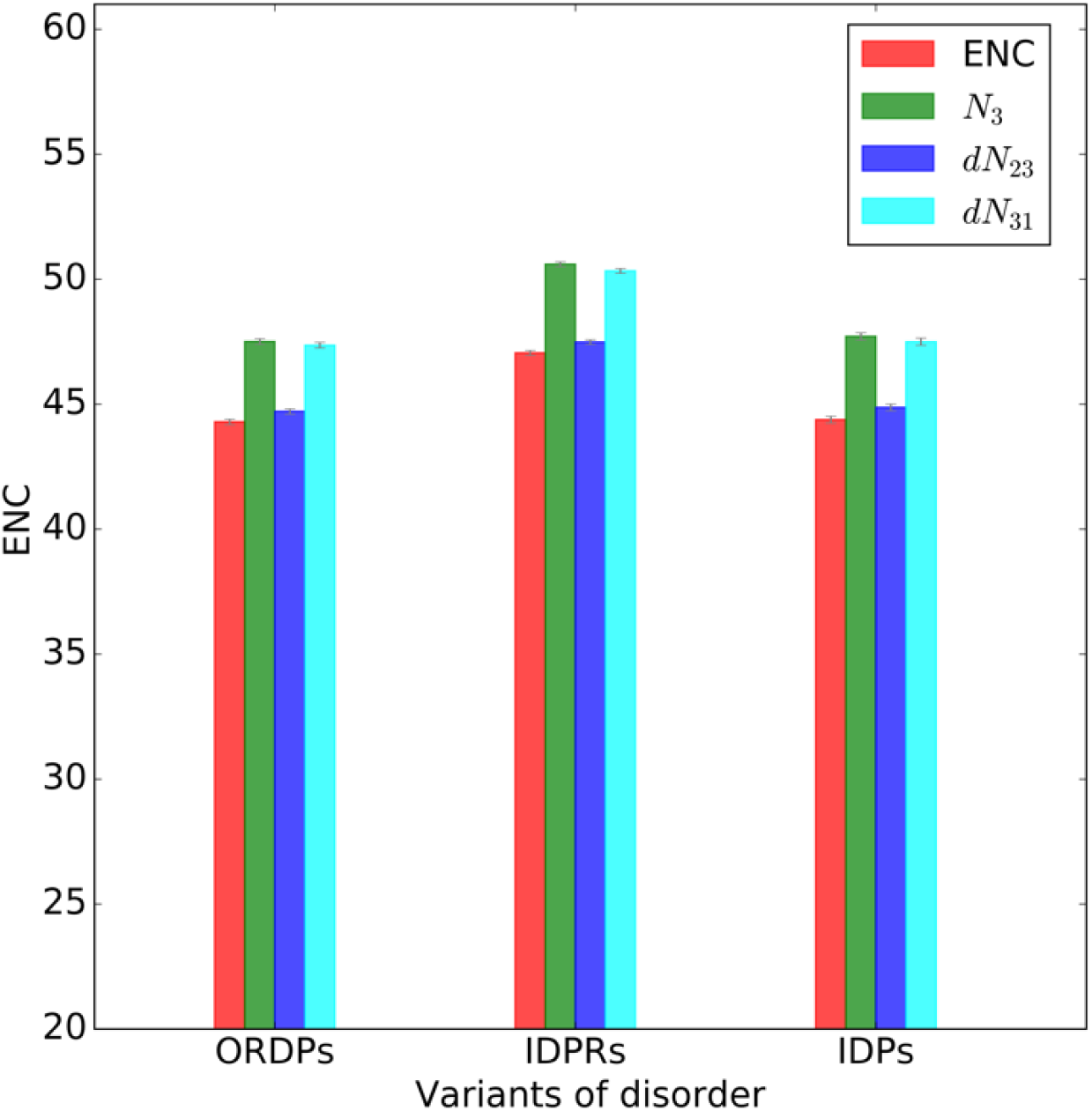
Average ENC values for the real coding sequences (in red) and shuffled sequences (other colors) obtained separately for all variants of disorder (ORDPs, IDPRs, and IDPs). On the x-axes, we show the variant of disorder; in the legend, we report the shuffling techniques used to analyze the impact of (di)nucleotide composition on CUB of the variants.

Secondly, we investigated the impact of dinucleotide content in codon positions 2-3 and 3-1 on CUB of the variants. Dinucleotide content in codon positions 1–2 was not considered in the present analysis being strongly constrained by the amino acid sequence. In all variants of proteins, shuffling of codon position 2-3 dinucleotide (dN_23_) produced ENC values very close to the original one, indicating a substantial impact of position 2-3 dinucleotide content on CUB of the variants (Fig. 7). Conversely, shuffling of dinucleotide position 3-1 produced values of ENC very similar to those obtained with the N_3_ correction (Fig. 7).

The differences between the ENC values obtained for the shuffled sequences and the corresponding values for the real coding sequences allow quantifying the extent of the unexplained residual bias (i.e., not explained by (di)nucleotide compositional bias). In Fig. 8 we report the average values of the differences obtained for each shuffling techniques in ORDPs, IDPRs, and IDPs. In this plot, closer to 0.0 is the average value of the difference and more we are justified in saying that the (di)nucleotide compositional bias used as a constraint for the shuffling algorithm is the cause of CUB. Clearly, most of CUB can be attributed to the compositional dinucleotide bias in codon position 2-3, but it still leaves a notable unexplained bias (insert on the bottom-right in Fig. 8). Notably, IDPs appear to be subject to a slightly stronger unexplained residual bias than ORDPs and IDPRs. Although the differences observed between IDPs and the other variants of disorder in terms of this unexplained residual bias are small, they are statistically significant (Welch’s t-test with Bonferroni correction for multiple comparisons, see insert on the right in Fig.8).

**Fig. 8:**
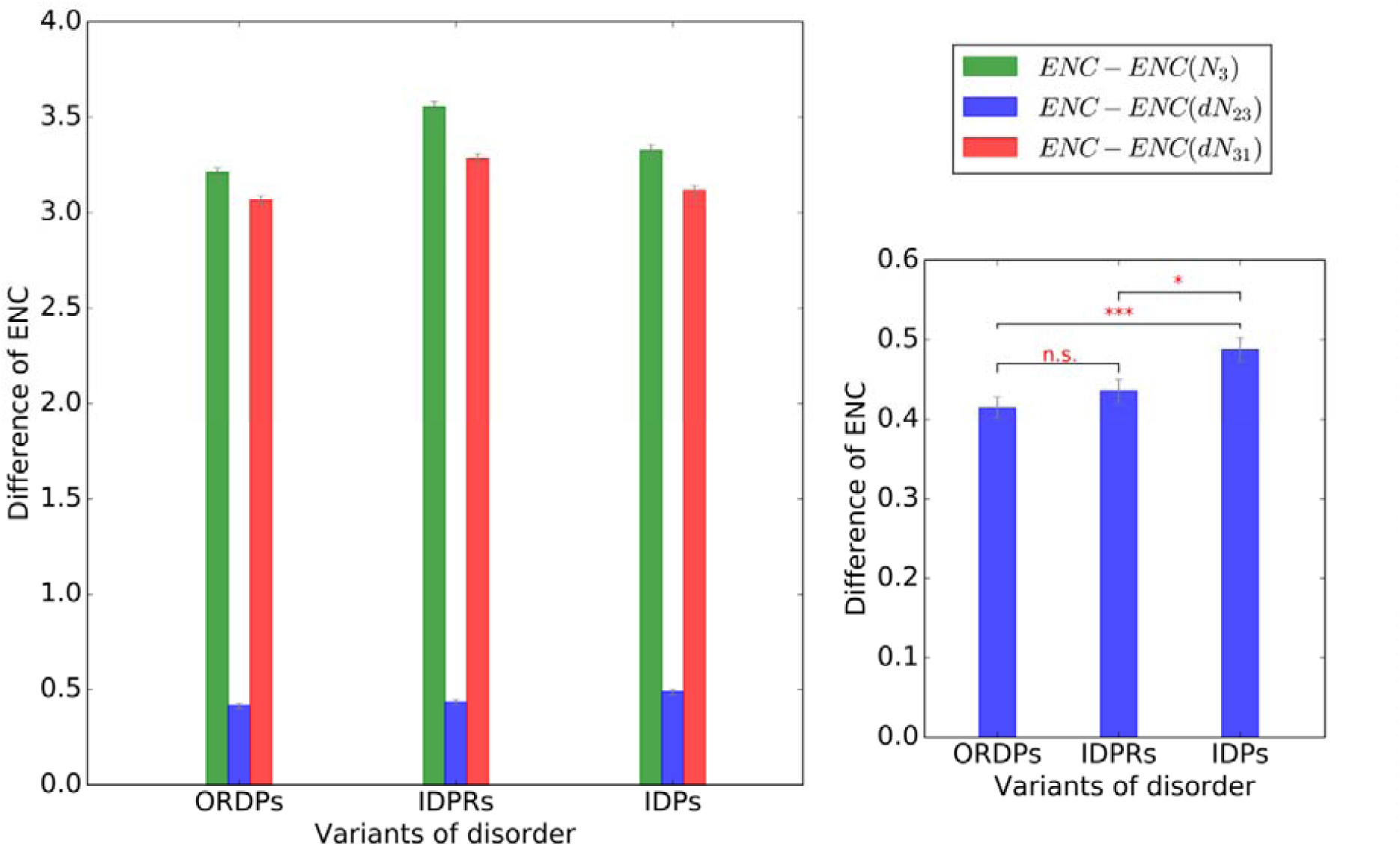
Average differences between ENC values obtained for N3, N23, and N31 shuffled sequences and the corresponding ENC values of natural coding sequences. On the bottom-right we report the average differences in ENC values obtained for N23 shuffled sequences and those for the original coding sequences. Significance is calculated using Welch’s t-test with adjustment for multiple testing. (***) < 0.001, (*) < 0.05, (n.s.) > 0.05.

### 3.6 Fraction of CpG sites in the coding sequences of variants of disorder

CpG sites are mutational hotspot in a wide range of human genes (Cooper, 1999). The high rate of C-T transition in methylated CpG sites represents an additional factor that strongly contributes to mutational pressure in the human genome (Bestor and Coxon, 1993; Pfeifer, 2006), and in shaping CUB of human genes. We, therefore, evaluated the abundances of CpG sites in ORDPs, IDPRs, and IDPs relative to what would be expected from the nucleotide composition of the variants (see Materials and Methods). CpG sites are overrepresented in IDPRs and IDPs and underrepresented in ORDPs (Fig. 9). Interestingly, IDPs are characterized by the highest fraction of CpG sites in the sequences, implying a higher susceptibility to methylation resulting in C-T transition mutations for IDPs than for IDPRs and ORDPs. The differences in CpG abundances were highly significance (Welch’s t-test with Bonferroni correction for multiple comparisons,).

**Fig. 9:**
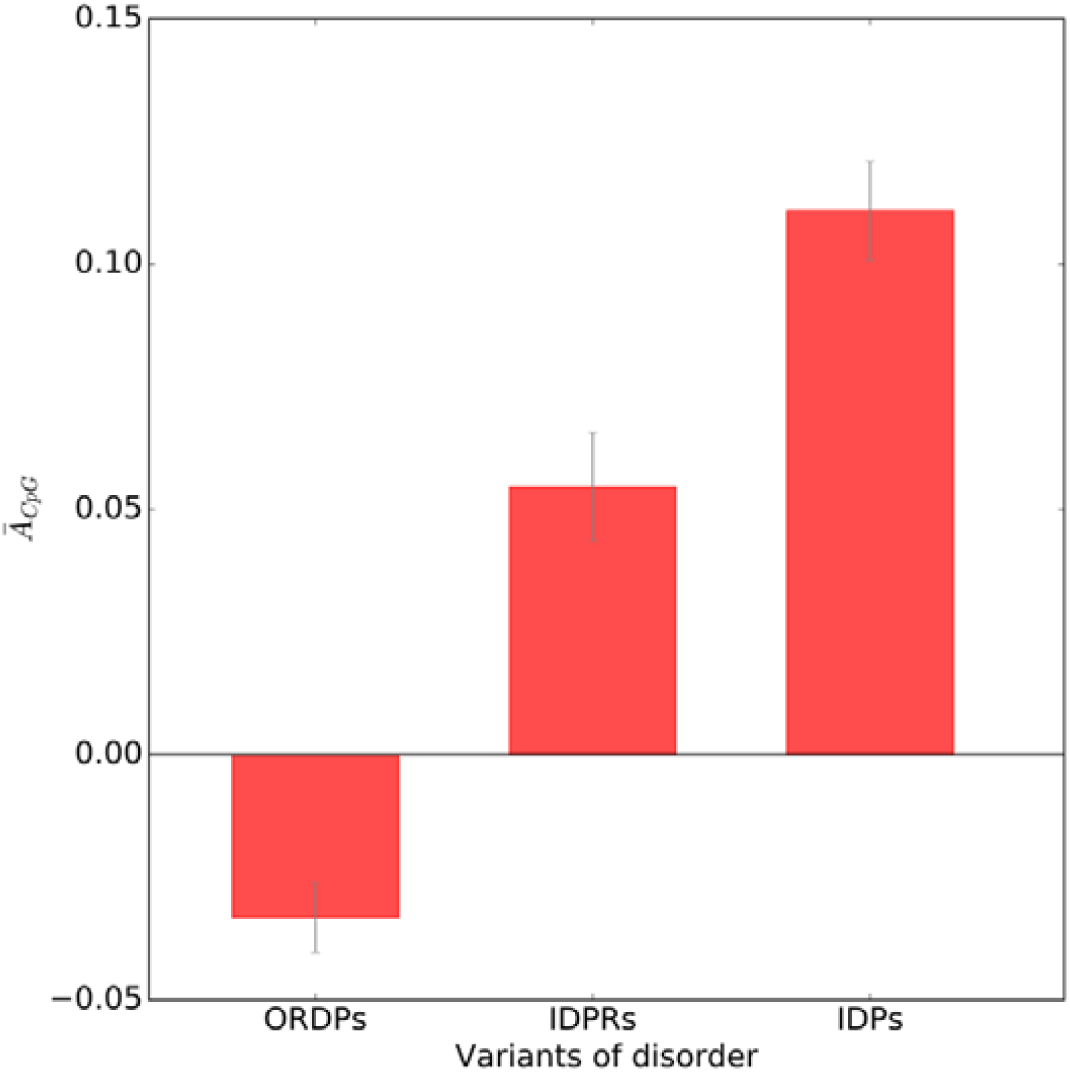
Average abundances of CpG-sites in human genes encoding for ORDPs, IDPRs, and IDPs with respect to what expected from the base composition of the variants and the standard errors on their assessments ().

## 4. Discussion

Analysis of protein evolution is central for many fields of research, including molecular evolution, comparative genomics, and structural biology (Pal et al., 2006). In this context, the phenomenon of codon usage bias (CUB) has been important because it allows quantifying both individual and combined effects of natural selection and mutational bias on genes.

With this study, we performed a systematic analysis of the evolutionary pressures (i.e., natural selection and mutational bias) that shape CUB of human genes encoding for different structural variants of proteins.

In line with a previous study (Deiana et al., 2019), we separated the human proteome in three broad variants of proteins characterized by different structural and functional properties: i) ordered proteins (ORDPs), ii) mostly ordered proteins with long intrinsically disordered protein regions (IDPRs), and iii) intrinsically disordered proteins (IDPs). ORDPs are expected to be more under control by natural selection than IDPs because one or few mutations (even synonymous) in the genes can result in a protein that no longer folds correctly. On the contrary, IDPs are generally thought to evolve more rapidly than well-structured proteins, primarily attributed to relaxed purifying selection due to the lack of structural constraints (Brown et al., 2002; Brown et al., 2010; Brown et al., 2011; Schlessinger et al., 2011; Xue et al., 2013). Suffice to say that specific subclasses of unstructured regions, such as entropic chains, may be under only two very general restrictions: keeping the region disordered and its length unchanged.

Using different genetic tools (such as ENC-plot, PR2-plot, and neutrality plot), we investigated the relative contribution of natural selection and mutational bias in shaping the base composition and CUB of genes encoding for proteins in ORDPs, IDPRs, and IDPs.

ENC-plot and PR2-plot of ORDPs, IDPRs, and IPDs show that both mutational bias and natural selection affect at different extent the CUB of all the variants of human proteins which is also supported by neutrality plot analysis (Fig. 1, Fig. 3). Looking at the best-fit curves in the ENC-plot and the positions of centroids in the PR2-plot, ORDPs and IPDs appear to be subjected on average to a similar balance between natural selection and mutational bias. On the contrary, IDPRs are the variant on which the contribution of mutational bias is more pronounced. This result strongly supports our previous claim to consider IDPRs and IDPs as distinct protein variants (Deiana et al., 2019), not only in terms of functional properties and roles in diseases but also in terms of evolutionary forces they are subjected to.

The neutrality plot analysis indicates that the first and second codon positions are slightly more neutral in IDPs than those in ORDPs and IDPRs (Fig. 4). This means that IDPs (i.e., mostly disordered proteins) are slightly freer to accept both synonymous and non-synonymous mutations without compromising their functionality in the cell.

However, it is worth noticing that none of the previous analyses considered any correction for background nucleotide or dinucleotide compositions, making it impossible to provide a direct estimation of the action of natural selection on CUB of the variants. For this purpose, we performed different shuffling techniques of the real coding sequences to filter all effects on CUB due to the background (di)nucleotide composition resulting from mutational processes. We conclude that most of CUB in all variants of proteins was attributed to mutational pressure on dinucleotide content in 2-3 codon position. However a weak (as expected by population genetic studies (Chamary et al., 2006)) but significant unexplained residual bias (i.e., not explained by background compositional bias) is revealed. In particular, IDPs are characterized by the highest extent of unexplainable bias (Fig. 8) that we here would identify with some selective mechanisms that differentiate them from the rest of human proteome.

Taken together, all these observations show that, despite the first and second codon positions are more neutral and thus more subjected to the mutational bias in IDPs than in ORDPs and IDPRs, the pattern of codon usage in IDPs is slightly more selective constrained than that of other variants of human proteins.

In line with Peng et al. (Peng et al., 2016), we reveal a strong positive correlation between the GC-content of genes and the level of intrinsic disorder in their corresponding proteins (Fig. 5, Fig. S1). We conclude that proteins with a high percentage of disordered residues (i.e., IDPs) are preferentially encoded by genes with high GC-content. Although genomic GC-content variations are commonly associated with the effect of mutational bias (Wolfe et al., 1989; Chen et al., 2004; Hershberg and Petrov, 2008; Plotkin and Kudla, 2011; Romiguier and Roux, 2017), several lines of evidence support a role of natural selection in maintaining high GC regions (Eyre-Walker and Hurst, 2001; Kudla et al., 2006; Li, 2013). In particular, it is well known that GC-rich genomic regions exhibit higher gene densities (International Human Genome Sequencing Consortium, 2001; Lercher et al., 2003; Versteeg et al., 2003), and are typically rich in highly expressed genes (Versteeg et al., 2003) and several biological activities (e.g., translation, transcription, and recombination) (Eyre-Walker and Hurst, 2001; Li, 2013). Moreover, it has been reported that housekeeping genes (in contrast to tissue-specific genes) are preferentially located in regions of high GC and, specifically, in CpG islands (Lercher et al., 2003). Because in human the GC-content of genes is related to the their genome location (Bernardi et al., 1985; Clay et al., 1996), all these considerations suggest a role of natural selection in controlling the long-time evolution of IDPs. Future studies could be aimed at investigating the fraction of human housekeeping genes that are also predicted as IDPs and the level of expression of this variant of human proteins, revisiting previous works (Lercher et al., 2003; Ma et al., 2014).

Another important factor that strongly contributes to mutational pressure in the human genomes is CpG methylation followed by deamination resulting in C-T substitution (Bestor and Coxon, 1993). C-T transition in CpG sites to form TpG dinucleotide gives rise to an underrepresentation of CpG dinucleotide (Simmen, 2008) that might strongly influence CUB of the genes. According to the high GC-content of genes encoding for IDPs, we observe that this variant of proteins has the highest fraction of CpG sites in the sequences, thus being potentially the variant more susceptible to C-T transition mutation (Fig. 9).

Methylation of the cytosine at CpG sites has important roles within the cell in regulating gene expression and silencing (Long et al., 2017). Single-gene and genome sequencing studies have identified that the vast majority of substitution mutations in human cancer occurs at the CpG dinucleotide (Cooper and Youssoufian, 1988; Esteller et al., 2001; Esteller, 2002; Poulos et al., 2017; Pfeifer, 2017). Genome-scale studies revealed that the most frequently methylated genes in cancer cells were genes encoding transcription factors (Pfeifer, 2018), which are specifically enriched in IDPs (Deiana et al., 2019). It has been claimed that about 30% of databases concerning mutations of tumor proteins p53 (here classified as IDP) is associated with transitions in CpG sites (Rodin and Rodin, 1998). Moreover, five major P53 mutational hot spots contain methylated CpGs (Denissenko et al., 1997). With this study we corroborate the observation that IDPs are over-represented in cancer-related proteins (Deiana et al., 2019) providing new insights about the role of this variant of proteins in carcinogenicity.

Many computational studies have shown that protein disorder is higher in eukaryotes relative to the less complex organisms (e.g., prokaryotes and archaea) (Dunker et al., 2000; Ward et al., 2004; Niklas et al., 2018). Additionally, it has been also revealed that high recombination and chromosomal rearrangement rates favor disordered regions during evolution, and relationships between GC-content, GC-biased gene conversion, and protein disorder have been established (Peng et al., 2016; Niklas et al., 2018). Such evidence suggests that intrinsically disordered proteins are essential for the evolution of new protein functions in eukaryotes and especially multicellular organisms (Schlessinger et al., 2011; Peng et al., 2016; Niklas et al., 2018).

In this study, we find evidence that IDPs are the variant of human proteins on which the main drivers of evolutionary change (i.e., mutational bias and natural selection) act more effectively, corroborating their hypothesized important role for evolutionary adaptability and protein evolvability (Tokuriki and Tawfik, 2009). At the same time, we speculate that IDPs have a high tolerance to mutations (both neutral and adaptive) but also a selective propensity to preserve their structural disorder, i.e., flexibility and conformational dynamics under physiological conditions. In support of that, different experimental and computational analyses showed that function and dynamic behavior of intrinsically disordered proteins are under selection and they are conserved in the face of negligible sequence conservation (Daughdrill et al., 2007; Lemas et al., 2016; Zarin et al., 2017; Walter et al., 2019). Additionally, there could be a strong selection to reduce the adverse effects of large concentrations of IDPs by tightly controlling their expression at all levels of transcription, translation, and protein degradation (Gsponer et al., 2008).

## 5. Conclusion

In this study, we performed a systematic analysis of the evolutionary pressures (natural selection and mutational bias) that shape synonymous codon usage patterns of human genes encoding for different structural variants of proteins. The main result is that intrinsically disordered proteins are not only affected by a basic mutational bias, but they also show a peculiar influence of natural selection as compared to the rest of human proteome. We find new evidence about the crucial role of intrinsically disordered proteins for evolutionary adaptability and protein evolvability identifying them as the variant of human proteins on which both mutational bias and natural selection act more effectively. Additionally, we confirm not only that intrinsically disordered proteins are preferentially encoded by GC-rich genes, but also that they are characterized by the highest fraction of CpG-sites in the sequences, implying a higher susceptibility to methylation resulting in C-T transition mutations (involved in several human cancers). Overall, our results provide new insight towards general laws of protein evolution identifying the intrinsically disordered proteins as reservoirs for evolutionary innovations. Quantifying selective and mutational forces acting on human genes could be useful for future studies concerning protein de novo design, synthetic biology, biotechnology applications, and the identification of proteins that are relevant in human genetic diseases. Moreover, extending the study presented here to the eukaryotes could add further comparative insight into the repertoire of the adaptive evolutionary mechanisms of proteins.

