## Supplementary material for "Evolutionary forces on different flavors of intrinsic disorder in the human proteome": FIgure S1.

Supplementary Informations

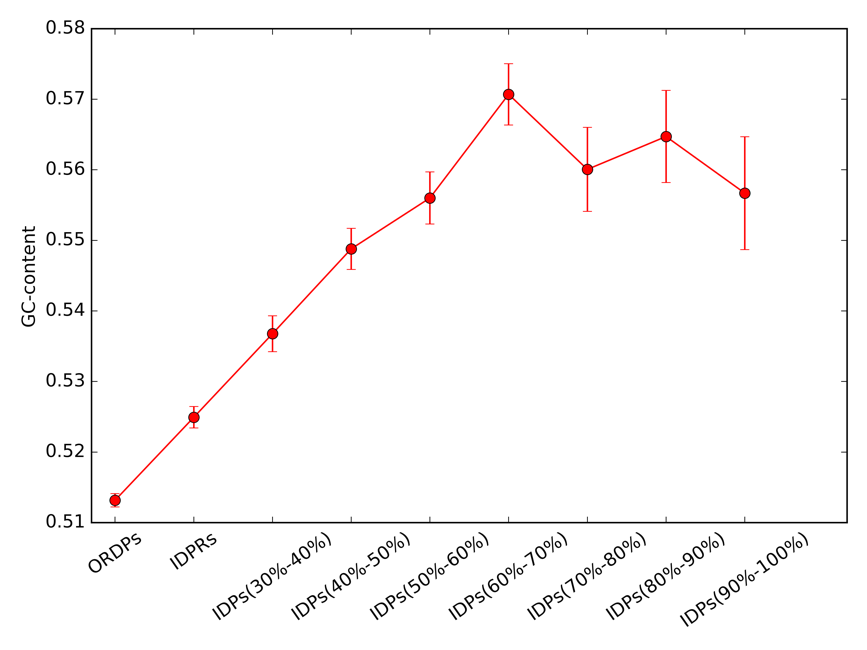

**Figure S1.** GC-content for ORDPs and IDPRs and for different percentiles of disordered residues in IDPs.
